# Individual-level Functional Connectivity Predicts Cognitive Control Efficiency

**DOI:** 10.1101/2022.07.14.500048

**Authors:** Benjamin L. Deck, Apoorva Kelkar, Brian Erickson, Fareshte Erani, Eric McConathey, Daniela Sacchetti, Olu Faseyitan, Roy Hamilton, John D. Medaglia

## Abstract

Cognitive control (CC) is a vital component of cognition associated with problem-solving in everyday life. Many neurological and neuropsychiatric conditions have deficits associated with CC. CC is composed of multiple behaviors including switching, inhibiting, and updating. The fronto-parietal control network B (FPCN-B), the dorsal attention network (DAN), the cingulo-opercular network (CON) and the dorsal default-mode network (dorsal-DMN) have been associated with switching and inhibiting behaviors. However, our understanding of how these brain regions interact to bring about CC behaviors is still unclear. In the current study, participants performed two in-scanner tasks that required switching and inhibiting. We then used a series of support vector regression (SVR) models containing individually-estimated functional connectivity between the networks of interest derived during tasks and at rest to predict inhibition and switching behaviors in individual subjects. We observed that the combination of between-network connectivity from these individually estimated functional networks predicted accurate and timely inhibition and switching behaviors in individuals. We also observed that the relationships between canonical task-positive and task-negative networks predicted inhibiting and switching behaviors. Finally, we observed a functional dissociation between the FPCN-A and FPCNB during rest, and task performance predicted inhibiting and switching behaviors. These results suggest that individually estimated networks can predict individual CC behaviors, that between-network functional connectivity estimated within individuals is vital to understanding how CC arises, and that the fractionation of the FPCN and the DMN may be associated with different behaviors than their canonically accepted behaviors.

## 1 Introduction

Cognitive control (CC) comprises processes that guide thoughts and actions to achieve goal-consistent actions, and is crucial to addressing the demands required in everyday tasks [1–3]. In psychological research, CC is conceptualized as three distinct yet related fundamental components of CC. These components are cognitive switching, inhibiting, and updating [4–7]. Our understanding of how brain regions interact to manifest CC is incomplete, as CC is reliant on circuits within the most topographically heterogeneous regions of the brain, specifically the prefrontal cortex (PFC) and lateral parietal lobes. Many studies of CC related networks examined brain-behavior associations using group-based atlases and short sequences of resting-state data [8], meaning that inferences about the role of brain networks in CC are confounded [5, 9–16]. Research suggests that individual differences in functional topology observed within PFC regions can predict an individual’s ability to deploy CC [17]. Furthermore, previous studies investigated individual differences in CC networks and provided information on behaviors mediated by spatially distinct brain regions in healthy and neuropsychiatric subjects [10, 12, 13, 16, 18–21].Despite these advances in understanding CC and the networks that subsume these behaviors, a gap in our knowledge remains about how these networks interact to inhibit responses and switch tasks in individuals. In the current study, we focus on four functional brain networks theorized to play critical roles in CC: the fronto-parietal control network (FPCN), the cingulo-opercular network (CON), the dorsal attention network (DAN), and the default-mode network (DMN) and investigate their relationship with one another to predict CC.

### 1.1 Cognitive functional brain networks associated with cognitive control

#### 1.1.1 The Fronto-parietal control network (FPCN)

The FPCN is frequently associated with switching and inhibiting responses and thoughts [22]. The FPCN interacts with the DMN, CON, and DAN during trial-level processing. Furthermore, the FPCN acts as a hub by fine-tuning sensory processing areas by biasing and integrating the areas’ information to achieve task goals [1–3, 10, 21, 23–40]. The functional heterogeneity of the FPCN complicates our understanding of its role regarding CC. Recent studies address the heterogeneity by dividing the FPCN into two networks: FPCNA and FPCN-B. Support for the division comes from research that showed activity within the FPCN-A was positively correlated with activity within the DMN during tasks that required internal attention. In contrast, activity within the FPCN-B correlated with the activity in the DAN during tasks that require external attention [10, 13, 16, 25, 30, 35, 41]. Previous work also showed that functional connectivity between the FPCN and other networks predicted an individual’s fluid intelligence [42]. This finding suggests that the relationships between functional networks may be meaningful to understanding an individual’s cognitive processes [43, 44]. However, previous studies have not investigated the variability in representation of FPCN-A, FPCN-B and associated activity between these two networks and other CC networks in individuals.

#### 1.1.2 The Cingulo-Opercular Network (CON)

The CON is active before, during, and after task stimuli are presented, which led researchers to conclude that the CON implements control by maintaining task goals and alertness [45, 46]. The CON is also vital for conflict and error monitoring after a stimulus is presented[31, 38, 47]. Furthermore, CON plays a role in maintaining alertness to given task goals. The FPCN likely interacts with CON and provides trial-by-trial level task decoding while CON maintains task-set goals crucial for successful completion of a task [48]. Previous studies suggest that these two networks work together to perform goal-directed behaviors where CON supports alertness and attention throughout a task[49]. If this model of task performance is accurate, then resting-state and task-based blood-oxygen-level-dependent (BOLD) connectivity between these two networks should predict overall task performance on tasks requiring continuous working memory and error monitoring updating.

#### 1.1.3 The Dorsal Attention Network (DAN)

The DAN triggers CC by shifting overt and covert spatial attention. Additionally, the DAN can bias top-down cognition and assist in selecting a response [50]. Specifically, the DAN orients a participant to task stimuli and helps them process sensory information [33, 50–52] during saccadic eye movements [53] and when a subject reaches for an essential object to complete a task [54]. Since the DAN is associated with spatial attention selection, DAN likely interacts with other CC networks, such as the FPCN, during cognitive tasks and is thought to provide essential sensory information to the FPCN for trial-to-trial manipulations of control [55].

#### 1.1.4 The Default Mode Network (DMN)

Activity within the DMN is frequently associated with the brain at rest (i.e., not participating in a goal-directed task) [56–58]. However, recent studies revealed that the DMN was active during other functions, which included introspective thinking [59], spontaneous cognition [60], working memory processing [61], creative problem solving/ divergent thinking [25], semantic memory processing [62], and task switching [34, 63]. Like the FPCN, the DMN also contains multiple distinct networks including: a midline network (canonical DMN behavior, i.e., mind-wandering) and a lateral component (associated with externally mediated tasks requiring internal mentation (i.e., task switching) [34, 56, 57, 62–67]). Interestingly, the FPCN can also causally influence activity within the DMN. In one study, the authors showed that transcranial magnetic stimulation (TMS) targeting the left-DLPFC (a node of the FPCN) caused a significant decrease in within-network connectivity in the DMN [68– 70]. Furthermore, previous work has also suggested that relationships between the FPCN, DAN, and DMN mediated better performance on working memory tasks in individuals with mild cognitive impairment, suggesting that normal working memory functioning may depend on how these networks co-vary [71]. While previous work suggests a strong anti-correlation between the canonical DMN and the FPCN, DAN, and CON is related to better CC task performance, to our knowledge these studies have not investigated the relationship between the FPCN subnetworks and dorsal-DMN [34]. Investigating how these networks are related to one another at rest and when performing CC tasks in individuals may help researchers to understand the complex functional diversity of the DMN and the FPCN.

### 1.2 The rationale for the current study

While previous studies demonstrated the functional independence of the FPCN, CON, DAN, and DMN and as such, that they likely support separate subprocesses related to CC [72], researchers have not yet examined how individually estimated functional connectivity between these networks is essential or predicts CC behaviors. The current study examines how individually estimated, resting-state, and task-based functional inter-network connectivity predicts task rule set switching (switch costs) and inhibition of prepotent behaviors (inhibition costs). We have four dependent hypotheses that we test in this study. **1)** We hypothesize that rest and task-based inter-network connectivity of FPCN-B, CON, and DAN predicts inhibiting and switching performance in single subjects. This hypothesis builds upon prior work indicating that each network mediates distinct subfunctions of CC: FPCN-B implements working memory updating of task sets and inhibits cued responses, CON maintains task goals and monitors for error responses, and DAN biases attention towards the target stimulus. **2)** Previous work suggests that the dorsal DMN supports task-set switching and working memory behavior. Therefore we hypothesize that connectivity between FPCN-B, DAN, CON, and dorsal-DMN will also predict inhibiting and switching in single subjects as the relationships between these networks will also support CC. **3)** Our third hypothesis is that the functions of FPCN-A and the B will dissociate, where FPCN-A will act more as a task-negative network (i.e., activity not involved in task performance) and will not meaningfully predict switching or inhibiting behaviors. **4)** Our final hypothesis consists of three sub hypotheses that examine how CC networks relate to one another. ***a)*** Task-based connectivity between FPCN-B and dorsal-DMN will positively correlate and negatively correlate at rest, ***b)*** Functional connectivity between FPCN-B, CON, and DAN will positively correlate during rest and task performance, and ***c)*** dorsal-DMN provides task-relevant information (positively correlated with CC networks) but couples negatively with CC networks at rest. See Table 2. Establishing a relationship between inter-network dynamics and cognitive control behaviors is crucial to understanding how networks support CC function at the level of the individual. Furthermore, these findings may guide personalized therapeutic advances that influence the entire distributed cognitive control system.

## 2 Methods

### 2.1 Participants

This study enrolled forty-two healthy right-handed adults (54.8% female), with a mean age of 26.6 (SD ± 7.53) years and with 17.05 (± 2.6) years of education. 59.5% of the forty-two subjects self-identified as white, 14.3%, self-identified as Black, 14.3% self-identified as Asian, 7% self-identified as multiple races, and 2.3% did not disclose. 88.1% self-identified as non-Hispanic, 2.3% self-identified as Hispanic, 2.3% self-identified as Hispanic or Latino, and a final 2.3% did not disclose. Participants were excluded if they had a history of neurological or psychiatric illness. All subjects volunteered with informed consent in writing under a protocol approved by the Institutional Review Board/ Human Subjects Committee at the University of Pennsylvania.

### 2.2 Behavioral Tasks

Participants performed either the Navon switching task, Stroop inhibition task, or both (See Figure 1). Participants were initially randomly assigned either the Navon or Stroop task. If they were interested in participating in a second session at least one week later, they performed the alternative task. 26.2% (11/42) of participants performed both tasks. All other subjects who enrolled only participated in one task. Altogether, twenty-nine subjects performed the Navon switching task, while twenty-one subjects performed the Stroop inhibition task.

**Fig. 1.**
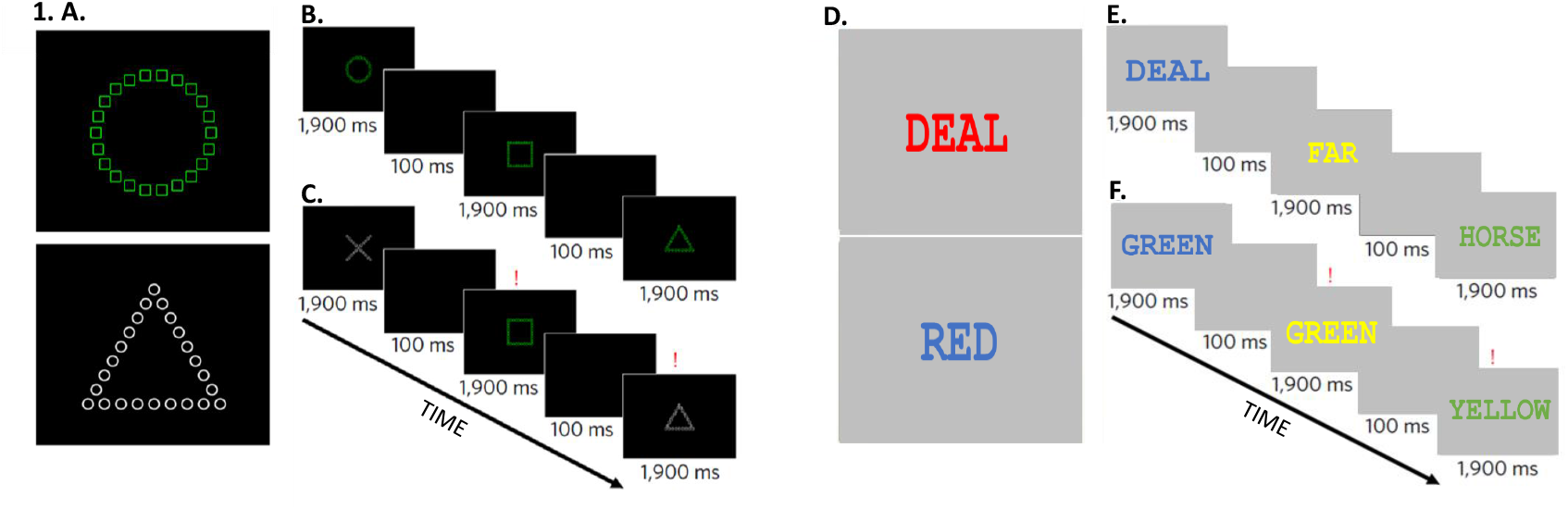
a) Example of global foreground stimulus indicating a circle is a correct response. Example of a local stimulus indicating a circle is the correct response b) Low difficulty trials consisting of global stimulus presented for 1,900 ms., followed by a blank screen presented for 100 ms. c) High-difficulty trials that require the subject to switch (indicated by a red exclamation point) between global and local stimuli. d) Example of low trial stimulus where non-relevant words are paired with colors. Example of a high trial stimulus where words are presented in incongruent ink colors. e) Example of a low difficulty trials, non-relevant words are presented in the target color ink for 1,900 ms., and a blank screen is presented for 100 ms. f) High difficulty trials which require the subject to inhibit (indicated by trials with a red exclamation point) the written response and respond to the color the ink is written.

The Navon switching task was adapted from Navon figures and required the subject to switch their perception from global to local fields or vice-versa [73, 74]. Briefly, the local-global stimuli consisted of four shapes – circle, X, triangle, or square. In all trials, the local and global features did not match. Stimuli were presented upon a black background in a block design with two block types. In block one, subjects saw white or green local-global stimuli. In block two, stimuli switched between white and green across trials uniformly at random, where 70% of all trials included a switch. If the stimuli were white, subjects were instructed to attend to only the local features. If the stimuli were green, subjects were instructed to attend to only the global features. Blocks were displayed in a random order per individual subject. Subjects responded using a four-button box corresponding to a circle, X, triangle, and square within the scanner. Subjects were trained on the task outside the scanner preceding the scan session. Each subject completed a set of practice trials for high and low conditions before completing the test block. There were 216 trials per test block. Each block was separated by a 30s fixation cross-hair presented in the center of the screen. Each trial was presented for 1900 ms, separated by a non-jittered inter-trial interval of a 100 ms black screen (Figure 1). We computed Navon switch-costs by subtracting the median low-trial responses from the median high-trial responses for response time and accuracy. The final switch-cost time was the time it took the subject to switch from local or global level stimuli or the accuracy trade-off.

The Stroop inhibition task requires subjects to inhibit automated responses [75]. This task consisted of high and low conditions. The high condition had trials with colored words corresponding to either red, yellow, green, or blue, printed in either matching or conflicting colored ink. Low condition trials included non-color words to control lexical demands without color-word conflict. The neutral terms were length and frequency matched to conflict trials. Responses were made using an in-scanner four-button box with red, yellow, green, and blue as possible responses. Font color was part of the target response set for all possible conditions. We calculated Stroop inhibition-costs by subtracting the median low trials from the median high trials for response time and accuracy.

### 2.3 Anatomical and functional image acquisition and preprocessing

#### 2.3.1 Acquisition

Anatomical images were acquired using T1w MP Rage volume (TR=2,400 ms; TE = 2.22 ms; flip angle = 8°; FOV = 256 mm; resolution 0.80 mm X 0.80 mm X 0.80 mm) and functional magnetic resonance imaging (fMRI) were acquired using a 3.0 Tesla Siemens Tim Trio whole-body scanner using a whole-head elliptical coil and a single-shot gradient-echo T2* (TR=500 ms; TE = 25.00 ms; flip angle = 30°; FOV = 192 mm; resolution 3.00 mm X 3.00 mm X 3.00 mm). Participants completed a resting-state scan lasting 8 minutes before completing either in-scanner Navon switching or Stroop inhibition tasks. Most subjects performed the Navon switching and Stroop inhibition tasks, and thus, many subjects had approximately 16 minutes of resting-state scan time.

#### 2.3.2 Anatomical data preprocessing

Anatomical preprocessing was performed using fMRIPrep 20.1.3 [76], based on Nipype 1.5.0 [77]. All T1-weighted (T1w) images were corrected for intensity non-uniformity (INU) with N4BiasFieldCorrection [78], distributed with ANTs 2.2.0 [79]. The T1w-reference was then skull-stripped with a Nipype implementation of the antsBrainExtraction workflow (from ANTs) using OASIS30ANTs as the target template [79]. Brain tissue segmentation of cerebrospinal fluid (CSF), white matter (WM), and gray matter (GM) were performed on the brain-extracted T1w using fast [80]. A T1w-reference map was computed after 3 T1w images (after INU-correction) registration using “mri robust template” [81]. Brain surfaces were reconstructed using recon-all [82]. The brain mask estimated previously was refined with a custom variation of the method to reconcile ANTs-derived and FreeSurfer-derived segmentation of the cortical gray-matter of Mindboggle [83]. Volume-based spatial normalization to one standard space (MNI152NLin2009cAsym) was performed through nonlinear registration with antsRegistration (ANTs 2.2.0), using brain-extracted versions of both T1w reference and the T1w template. The following template was selected for spatial normalization: ICBM 152 Nonlinear Asymmetrical template version 2009c [84]; TemplateFlow ID: MNI152NLin2009cAsym].

#### 2.3.3 Functional data preprocessing

For each BOLD run preprocessed per subject (across all tasks and sessions), a reference volume and its skull-stripped version were generated using a custom methodology of fMRIPrep [76]. Head-motion parameters for the BOLD reference (transformation matrices and six corresponding rotation and translation parameters) were estimated before any spatiotemporal filtering using mcflirt (FSL 5.0.9, [85]. BOLD runs were slice-time corrected using 3dTshift from AFNI 20160207 [86].

A B0-nonuniformity map (or fieldmap) was estimated based on a phase-difference map calculated with a dual-echo GRE (gradient-recall echo) sequence, processed with a custom workflow of SDCFlows inspired by the epidewarp.fsl script and further improvements in HCP Pipelines [87]. The fieldmap was then co-registered to the target EPI (echo-planar imaging) reference run and converted to a displacements field map (amenable to registration tools such as ANTs) with FSL’s fugue and other SDCflows tools. A corrected EPI (echo-planar imaging) reference was calculated for a more accurate coregistration with the anatomical reference based on the estimated susceptibility distortion.

The BOLD reference was then co-registered to the T1w reference using bbregister (FreeSurfer), which implements boundary-based registration [88]. Co-registration was configured with six degrees of freedom. The BOLD time-series were resampled onto the following surfaces (FreeSurfer reconstruction nomenclature): fsaverage. The BOLD time-series (including slice-timing correction when applied) were resampled onto their original, native space by applying a single, composite transform to correct for head-motion and susceptibility distortions. These resampled BOLD time-series were referred to as preprocessed BOLD in original space or just preprocessed BOLD.

Several confounding time-series were calculated based on the preprocessed BOLD: framewise displacement (FD), DVARS, and three region-wise global signals. FD was computed using two formulations following Power (absolute sum of relative motions [89] and Jenkinson (relative root mean square displacement between affines [85]). FD and DVARS were calculated for each functional run, using their implementations in Nipype (following the definitions by [89]). The three global signals were extracted within the CSF, the WM, and the whole-brain masks.

Additionally, a set of physiological regressors were extracted to allow component-based noise correction [90]). Principal components were estimated after high-pass filtering the preprocessed BOLD time-series (using a discrete cosine filter with 128s cut-off) for the two CompCor variants: temporal (tCompCor) and anatomical (aCompCor). tCompCor components are then calculated from the top 5% variable voxels within a mask covering the subcortical regions. This subcortical mask was obtained by heavily eroding the brain mask, which ensured it does not include cortical GM regions. For aCompCor, components were calculated within the intersection of the mask as mentioned earlier, and the union of CSF and WM masks was calculated in T1w space after their projection to the native space of each functional run (using the inverse BOLD-to-T1w transformation). Components were also calculated separately within the WM and CSF masks. For each CompCor decomposition, the k components with the largest singular values were retained, such that the retained components’ time series was sufficient to explain 50 percent of variance across the nuisance mask (CSF, WM, combined, or temporal). The remaining components were dropped from consideration.

The head-motion estimates calculated in the correction step were also placed within the corresponding confounds file. confound time series derived from head motion estimates and global signals were expanded to include temporal derivatives and quadratic terms for each [91]. Frames that exceeded a threshold of 0.5 mm FD or 1.5 standardized DVARS were annotated as motion outliers. All resampling was performed with a single interpolation step by composing all the pertinent transformation (i.e., head-motion transform matrices, susceptibility distortion correction when available, and co-registrations to anatomical and output spaces). Gridded (volumetric) resampling was performed using antsApplyTransforms (ANTs), configured with Lanczos interpolation to minimize the smoothing effects of other kernels [92]. Non-gridded (surface) resampling was performed using mri vol2surf (FreeSurfer). Many internal operations of fMRIPrep use Nilearn 0.6.2 [93], mainly within the functional processing workflow.

All functional data were preprocessed further using XCP engine (version 1.2.1) to perform confound regression and residualizing physiological artifacts from individual BOLD images [94]. XCP engine uses the output from fMRIprep and applies a modular structure to constructing a preprocessing pipeline. We implemented the fc-36p-despike design file for resting-state data (https://github.com/PennBBL/xcpEngine). The 36p-despike design file applies the following modules: prestats which generates several outputs derived from each subject’s fMRIprep run and runs image quality control measures; confound2, which models artifact signal in the 4D image time series including the white matter, grey matter, cerebral spinal fluid, and global signal; and the regress module which takes the output of confound2 module and fits a multiple linear regression on the time series and the explained variance in the BOLD time series from confounds are removed, and the residuals are used as the preprocessed time-series. Within the regress modules, despiking also occurred where large spikes within the BOLD time series were truncated.

We also implemented XCP engine (version 1.2.1) using the task design file to perform FSL feat twice to regress out artifact from the time series associated with task performance and a second iteration where non-task-related activity was regressed out of the time series, i.e., activity related to cortical motor activation due to button box responses [95].See overview of image preprocessing and estimation of functional networks in Figure 2

**Fig. 2.**
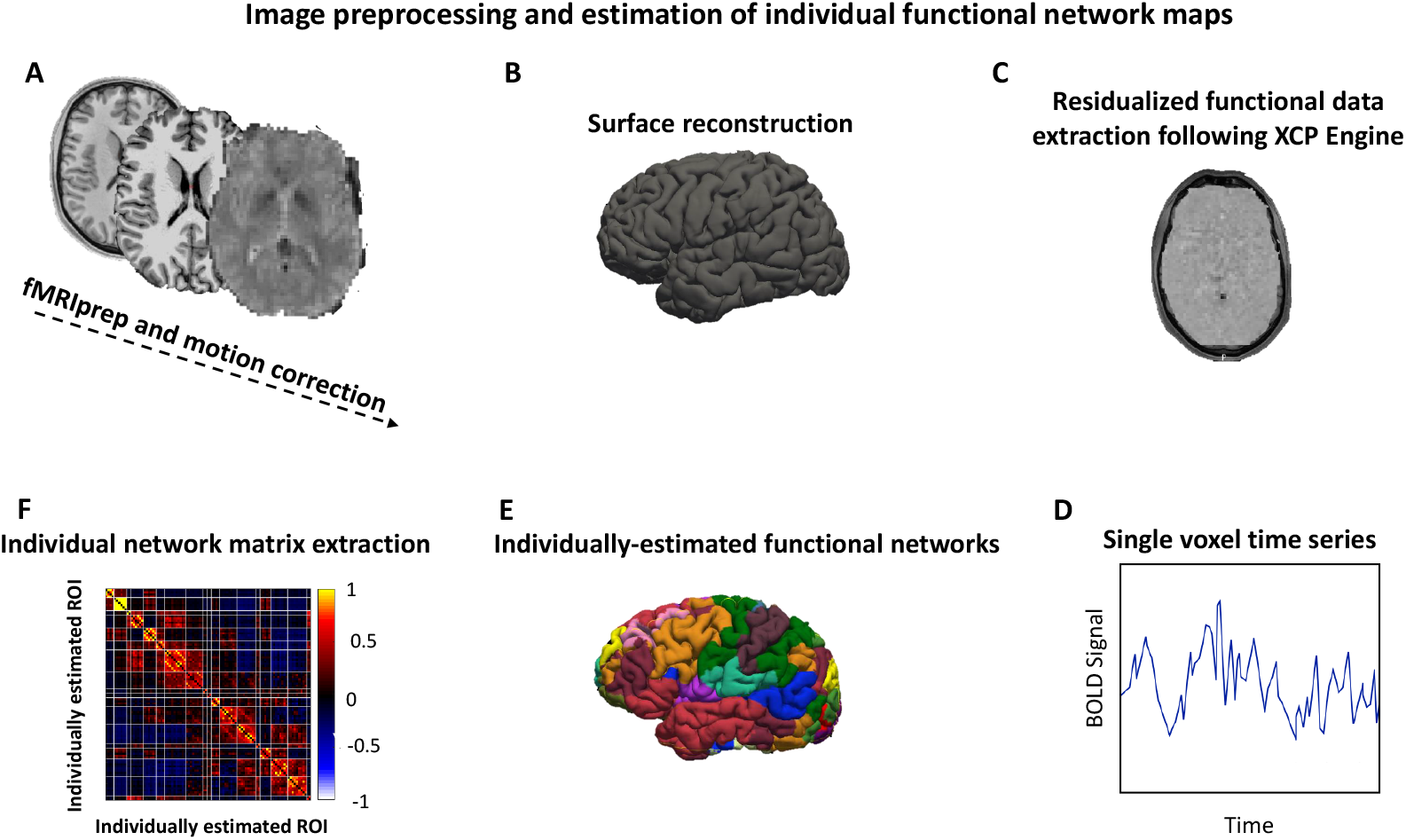
.A) Each subject’s T1w, rest, and task images were motion corrected via fMRI prep and XCP engine. B) All T1w images were registered to fsavearge template so that Surface reconstruction via Freesurfer Recon-all could be performed. C) Regression of functional time series using despiking protocol from XCP Engine. D) Representation of preprocessed BOLD time series from a single voxel used as input to Li et al., 2019 software. E) Individually estimated intrinsic networks were estimated in volumetric space and then re-registered to the individual subject surface for visualization and inspection. F) Z-scored intrinsic connectivity matrices were extracted for each individual. Further averaging across networks is not depicted here. See the Statistical Analysis section for details.

#### 2.3.4 Individualized Functional Network Mapping

Performing subject-level connectivity analysis accounts for inter-subject variability in functional topography and topology not otherwise accounted for using group atlases [13, 16, 20]. We performed connectivity mapping within individuals using a previously described methodology [13, 16]. Of the 116 total ROIs assumed by the parcellation, we found approximately 83.6% of all ROIs across all individuals during rest. We observed an average of 73.1% of all ROIs across all individuals during the Navon switching task and Stroop inhibition task performance. Limitations in the amount of resting-state and task data collected or missing functional ROIs in individuals may explain why we did not detect smaller ROIs [13, 16, 20]. However, the number of ROIs acquired was similar to previous studies examining individually mapped ROIs within healthy controls [13, 16]. If a specific ROI was not mapped to an individual, the ROI was dropped, and the region was identified according to the subsequent highest seed correlation. See Figure 2 for an overview of imaging methods.

#### 2.3.5 Statistical Analysis

Functional connectivity was extracted from the BOLD time-series using Li and colleague’s (2019) software [13]. The result was a pairwise ROI-by-ROI matrix per individual with Fisher’s exact *z* -scores derived from Pearson’s *r* correlation coefficients that made up the edges of the matrices [13, 96–98]. We extracted functional connectivity matrices from every individual during rest and task performance (high and low conditions separately for the Navon switching and Stroop inhibition tasks). We subtracted each individual’s functional connectivity matrix in the low task condition from the high condition for the task data. The high minus low contrast reflects connectivity changes due to switching task rule sets and inhibiting automated responses for the Navon and Stroop. Then, we computed the average connectivity value between each pair of regions between each network of interest (FPCN-A, FPCN-B, dorsal-DMN, CON, and DAN). Between-network connectivity in this study is the statistical relationship (Fisher’s exact z-score) between the networks of interest. Each average inter-network connectivity measure was used as an input feature. The result was a subject-by-feature matrix which consisted of rows of subjects, columns of between network connectivity from each pair of regions, and a target vector of either median response time or mean accuracy for the resting-state, Navon switching, and Stroop inhibition high (–) low contrast conditions.

Our feature set consisted of a series of columns. Each column was the average between network connectivity of our regions of interest (ROIs), namely, FPCN-B, FPCN-A, dorsal-DMN, CON, and DAN. When we examined the dissociation between FPCN-A and FPCN-B, we replaced all columns between FPCN-B and other networks with FPCN-A and all other networks.

To test our hypotheses that inter-network connectivity is related to CC behaviors, we used a support vector regression (SVR) technique. We justify utilizing this technique as SVR models can model high-dimensional nonlinear data. We implemented our SVR models with Scikit-learn in Python [99]. We chose the SVR model parameters via 10-fold cross-validation using a grid-search cross-validation. The grid-search cross-validation technique computes all possible models from a range of Cost (a regularization parameter that acts to penalize data that fit the model poorly), gamma (a kernel coefficient, which allows the model to fit across high dimensional data), epsilon (a margin of error before C is applied), and using one of the following kernels: radial base function, linear or sigmoid (see supplemental). Using a grid-search cross-validation technique, we determined the best fitting model given by a mean squared error (MSE). Our cross-validation technique also tested our model using different combinations of subgroups using an 80% to 20% split for training and test data, respectively, with 10x folds (i.e., ten folds).

We obtained an optimal mean-squared error value from our cross-validation process. We deployed a bootstrap method using the same model parameters as our accepted optimal model. We constructed a bootstrap method (10-thousand bootstrap samples) to develop a 95% confidence interval around the given MSE. The bootstrap method we deployed resampled 50% of the total subject count for each test, refitting the SVR model and obtaining a new MSE.

The same connectivity and behavioral data were submitted to a permutation test, which tested whether the observed MSE value exceeded that obtained from a random pairing between connectivity and behavioral data. We generated a null distribution of MSE values by performing 10,000 random permutations of the features (functional connectivity) and the outcomes (behavioral scales) and fitting an SVR model for each permutation. A p-value was obtained by calculating the proportion of MSE values in the permuted distribution that performed worse than the observed model fit (i.e., a higher MSE value). The model was deemed statistically significant if the model performed better than 95% of the null distribution.

We used one sample two-way t-tests to test whether there were significant correlations between the CC networks. We set the null hypothesis equal to 0, indicating that the networks were not correlated.

Correction for multiple comparisons was performed using the BenjaminiHochberg method of correction for multiple comparisons [100]. This correction set the overall study alpha threshold (alpha= 0.015) with an approximate false discovery rate (FDR) at 2.5% of the overall findings. Setting a threshold at an FDR of 2.5% meant that at 58 independent tests, approximately 1.45 of the tests could be a false positive and thus a Type I error (all statistical tests and their corresponding Benjamini-Hochberg FDR value are given in the supplement). Our final t-tests containing the FPCN-A network assessed whether there were significant correlations compared to 0 between the CC networks and were exploratory rather than confirmatory; we did not correct for multiple comparisons [101].

We used random forest regression models to perform a post-hoc feature analysis [102] to examine the relative importance of predicting switch-cost and inhibition-cost accuracy and response time. Random forest regression is an adaptation of decision tree regression that computes multiple decision trees and averages the trees together to determine an overall MSE that predicts a given set of targets or dependent variables [102]. One advantage of using random forest regression is that each sample of data points is drawn from the original sample with a replacement, i.e., a bootstrap sample. Each node split is based on a random subset of features. The advantage of using such levels of randomness is that it decreases the variance of the estimator of random forest models, in this case, the residual sum-of-squares [102, 103]. Singular forest models tend to have high variance and overfit; however, one of the tuning parameters is the number of forests to compute. Based on the overall best model fit for a set of features and targets, the forests were averaged, and this averaging process moderates the errors of single models. The hyperparameters were selected using a randomized search cross-validation method with one hundred different parameter sets per fold, which examined an extensive array of model parameters across ten cross-validation folds (see supplement for a list of possible parameters) [104]. Using random forest regression to determine relative levels of important features to predict the outcome variable allowed us to compute the importance of a feature relative to all other features in the model by considering their correlation. The feature importance test takes advantage of bootstrapping in random forest regression. The out-of-bag (OOB) or test sample examines the importance of predicting the target variable on a random selection of features [102, 103].

We also permuted each feature one thousand times and established a relative distribution of the importance of each feature using MSE. If the overall model fit is unchanged (i.e., MSE stays mostly the same), then the variable is given a lower value for its importance, i.e., closer to 0. In contrast, a feature that, when permuted, dramatically increases the overall model MSE would receive a higher value for importance, closer to 1. The calculation of importance is computed by averaging the difference in OOB before and after the permutation of all trees. The score is procured by using the standard deviation of the differences in OOB fits before and after the permutation of all trees to normalize the score [105].

## 3 Results

### 3.1 Hypothesis 1: Functional connectivity between FPCN-B, CON, and DAN predicts set switching and inhibition of prepotent responses in single subjects

We examined our first hypothesis that inter-network functional connectivity would predict switch-cost and inhibition-cost accuracy and response time. We observed partial support for our first hypothesis that resting-state connectivity between the FPCN-B, the DAN, and the CON significantly predicted median response time of switching and inhibition-costs. However, rest connectivity between the FPCN-B, DAN, and CON only significantly predicted mean accuracy of switch-costs better than null distributions (Switch-Cost RT: MSE-hypothesized model = 0.99, (Root-MSE = 0.98), 95% CI [0.62, 1.66], median MSE of permuted models = 2.01 (4.04), p = 0.004, Switch-Cost Accuracy: MSE-hypothesized model = 0.86 (0.74), 95% CI [0.70 1.73], median MSE of permuted models = 1.35 (1.82), p = 0.013; Inhibition-Cost RT: MSE-hypothesized model = 0.79 (0.62), 95% CI [0.45, 1.82], median MSE of permuted models = 2.00 (4.02), p = 0.002, Inhibition-Cost Accuracy: MSE-hypothesized model = 0.85 (0.72), 95% CI [0.60 3.89], median MSE of permuted models = 1.40 (4.04), p = 0.02). See Figure 3

**Fig. 3.**
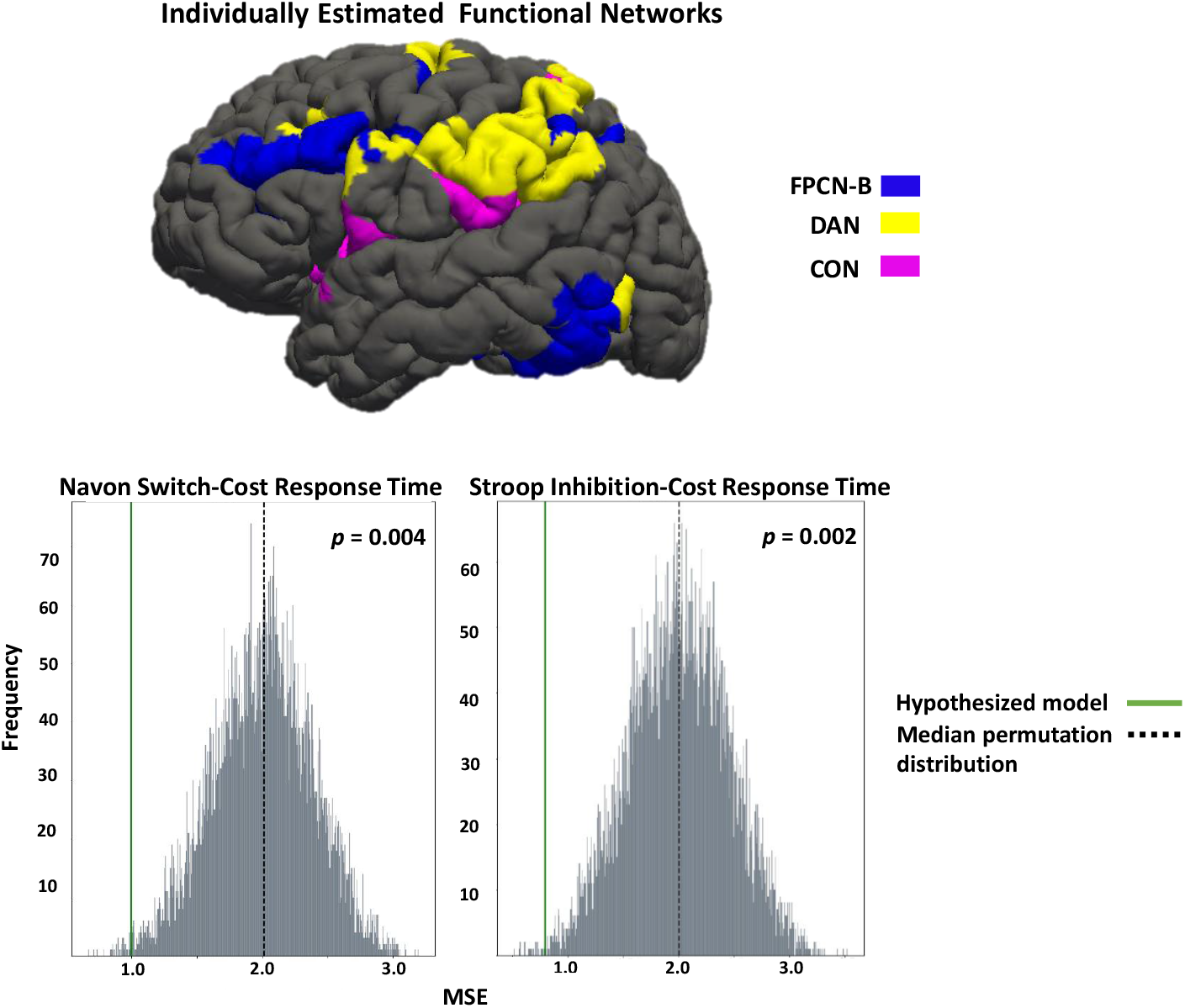
Top) Surface-based representation of networks that predicted switching and inhibiting behaviors in individuals. Bottom) Permutation tests comparing our hypothesized model (connectivity between FPCN-B, CON, and DAN predicting switching and inhibiting in individuals) to a random null distribution. Both performed better than random distributions at p = 0.004 and p = 0.002, respectively.

We observed further support for our first hypothesis by observing *taskbased* functional connectivity between the same three networks significantly predicted accuracy of switch-cost and inhibition-cost accuracy but only inhibition-cost response time better than null distributions (Switch-Cost Accuracy: MSE-hypothesized model = 0.98 (0.96), 95% CI [0.69 1.89], median MSE of permuted models = 1.00 (1.00), p = 0.0001; Inhibition-Cost Accuracy: MSE-hypothesized model = 0.88 (0.77), 95% CI [0.58 2.60], median MSE of permuted models = 1.12 (1.25), p = 0.009; Switch-Cost RT: MSE-hypothesized model = 1.02 (1.04), 95% CI [0.65 1.73], median MSE of permuted models = 1.48 (2.19), p = 0.04; Inhibition-Cost RT: MSE-hypothesized model = 0.58 (0.34), 95% CI [0.69 8.74], median MSE of permuted models = 1.56 (2.43), p = 0.003.

### 3.2 Hypothesis 2: Functional connectivity between FPCN-B, CON, DAN, and dorsal-DMN predicts set switching and inhibiting automated responses in single subjects

In partial support for our second hypothesis we observed that *resting-state* functional connectivity between FPCN-B, DAN, CON, and dorsal-DMN significantly predicted median response time and mean accuracy of switch-costs but not inhibition costs better than null distributions (Switch-Cost RT: MSE-hypothesized model = 0.99 (1.00), 95% CI [0.63 1.66], median MSE of permuted models = 1.99 (3.98), p = 0.005, Switch-Cost Accuracy: MSE-hypothesized model = 0.79 (0.62), 95% CI [0.60, 1.58], median MSE of permuted models = 2.00 (4.01), p = 0.0001; Inhibition-Cost RT: MSE-hypothesized model = 0.89 (1.78), 95% CI [0.62 2.17], median MSE of permuted models = 1.16 (1.35), p = 0.07, Inhibition-Cost Accuracy: MSE-hypothesized model = 1.05 (1.10), 95% CI [0.59 1.86], median MSE of permuted models = 1.00 (1.01), p = 0.94).)

In further support of this hypothesis we observed that the *task-based* internetwork connectivity of these networks predicted switch-cost and inhibition-cost accuracy but not response time better than null distributions (Switch-Cost Accuracy: (MSE-hypothesized model = 0.78 (0.60), 95% CI [0.68 1.94], median MSE of permuted models = 1.07 (1.14), p = 0.0002; Inhibition-Cost Accuracy: (MSE-hypothesized model = 0.60 (0.37), 95% CI [0.66 5.79], median MSE of permuted models = 1.44 (2.07), p = 0.005; Switch-Cost RT: MSE-hypothesized model = 1.05 (1.10), 95% CI [0.69 1.76], median MSE of permuted models = 1.01 (1.02), p = 1.00; Inhibition-Cost RT: MSE-hypothesized model = 0.99 (0.98), 95% CI [0.59 1.97], median MSE of permuted models = 1.01 (1.02), p = 0.36).

Within these omnibus models, we determined that the most important feature in predicting switch-cost response time was *resting-state* connectivity between FPCN-B and DAN. In contrast, *resting-state* connectivity between dorsal-DMN and DAN was most important in predicting switch-cost accuracy, inhibition-cost response time, and inhibition-cost accuracy. We again examined the relative feature importance in predicting inhibition and switch-cost response time and accuracy using inter-network connectivity during *task* performance. We observed that task-based inter-network connectivity between dorsal-DMN and DAN was the most important feature in predicting response time and accuracy for switch-costs (Table 1). Furthermore, we also observed that the *task-based* inter-network connectivity of FPCN-B and CON was the most important feature in predicting inhibition-cost response time and accuracy. See Table 2

**Table 1.**
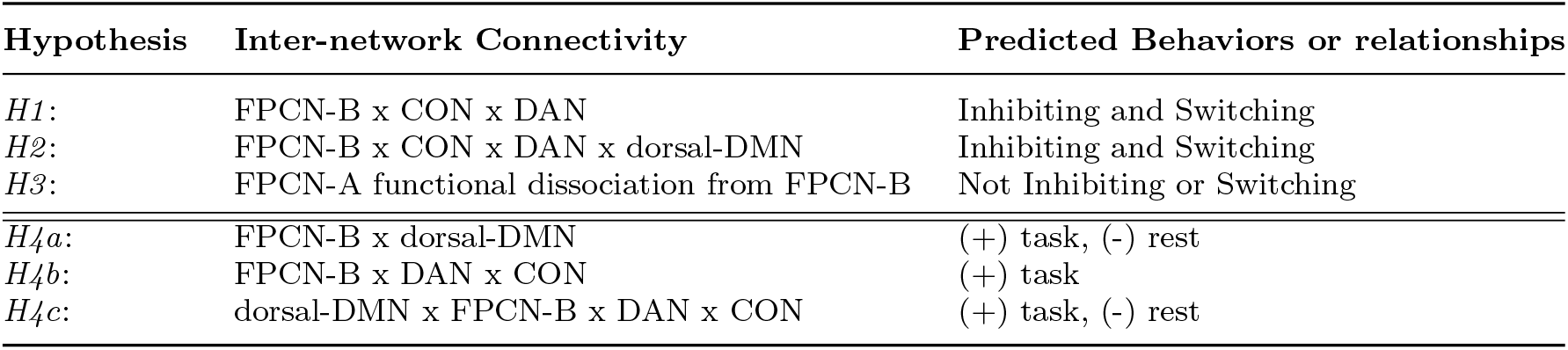
Table of Hypotheses

**Table 2.** * indicates the most important feature of an analysis

| Rest | Switch-cost Response Time (mean and std OOB) | Switch-cost accuracy |
| --- | --- | --- |
| FPCN-B x DAN | **0.11 (0.04)\*** | 0.04 (0.2) |
| FPCN-B x CON | 0.03 (0.01) | 0.04 (0.01) |
| CON x DAN | 0.09 (0.04) | 0.08 (0.03) |
| FPCN-B x d-DMN | 0.09 (0.04) | 0.09 (0.04) |
| d-DMN x CON | 0.06 (0.02) | 0.05 (0.02) |
| d-DMN x DAN | 0.04 (0.02) | **0.12 (0.05)\*** |
| Task |  |  |
| FPCN-B x DAN | 0.05 (0.02) | 0.01 (0.003) |
| FPCN-B x CON | 0.09 (0.03) | 0.06 (0.02) |
| CON x DAN | 0.05 (0.01) | 0.08 (0.03) |
| FPCN-B x d-DMN | 0.12 (0.04) | 0.02 (0.01) |
| d-DMN x CON | 0.04 (0.01) | 0.11 (0.03) |
| d-DMN x DAN | **0.21 (0.07)\*** | **0.28 (0.09)\*** |
| Rest | Inhibition-cost Response Time | Inhibition-cost accuracy |
| FPCN-B x DAN | 0.04 (0.02) | 0.04 (0.02) |
| FPCN-B x CON | 0.08 (0.02) | 0.02 (0.01) |
| CON x DAN | 0.07 (0.03) | 0.08 (0.04) |
| FPCN-B x d-DMN | 0.10 (0.04) | 0.03 (0.01) |
| d-DMN x CON | 0.16 (0.05) | 0.03 (0.01) |
| d-DMN x DAN | **0.31 (0.11)\*** | **0.15 (0.06)\*** |
| Task |  |  |
| FPCN-B x DAN | 0.09 (0.04) | 0.02 (0.01) |
| FPCN-B x CON | **0.17 (0.04)\*** | **0.28 (0.09)\*** |
| CON x DAN | 0.03 (0.02) | 0.11 (0.04) |
| FPCN-B x d-DMN | 0.04 (0.03) | 0.10 (0.05) |
| d-DMN x CON | 0.09 (0.04) | 0.15 (0.07) |
| d-DMN x DAN | 0.06 (0.03) | 0.10 (0.04) |

### 3.3 Hypothesis 3: Functional connectivity between FPCN-A and CC networks would predict dissociable behaviors in single subjects compared with functional connectivity between FPCN-B and CC networks

To test our third hypothesis that there is a dissociation between FPCN-A and FPCN-B, we examined the same models previously described with FPCN-A instead of FPCN-B. We observed partial support for our third hypothesis by observing that *resting-state* connectivity between FPCN-A, CON, and DAN significantly predicted mean switch-cost accuracy, but not mean inhibition-cost accuracy or median response time of switch-costs but did predict inhibition-cost response time better than null distributions (Switch-Cost Accuracy: MSE-hypothesized model = 0.87 (0.76), 95% CI [0.62 1.70], median MSE of permuted models = 1.05 (1.10), p = 0.006; Inhibition-Cost Accuracy: MSE-hypothesized model = 1.05 (1.10), 95% CI [0.60 1.88], median MSE of permuted models = 1.00 (1.00), p = 0.94; Switch-Cost RT: MSE-hypothesized model = 0.96 (0.92), 95% CI [0.63 2.00], median MSE of permuted models = 1.06 (1.12), p = 0.14; Inhibition-Cost RT: MSE-hypothesized model = 0.87 (0.76), 95% CI [0.66 1.91], median MSE of permuted models = 1.86 (3.45), p = 0.009).

Additionally, we also observed that *task-based* functional connectivity between these networks measured during the Navon switching task did not significantly predict accuracy or response times of switch-costs or inhibition-costs better than null distributions (Switch-Cost Accuracy: MSE-hypothesized model = 0.98 (0.96), 95% CI [0.69 1.75], median MSE of permuted models = 1.02 (1.04), p = 0.22; Switch-Cost RT: MSE-hypothesized model = 0.99 (0.98), 95% CI [0.70 1.81], median MSE of permuted models = 1.03 (1.06), p = 0.31; Inhibition-Costs Accuracy: MSE-hypothesized model = 0.83 (0.69), 95% CI [0.56 1.88], median MSE of permuted models = 1.34 (1.80), p = 0.03; Inhibition-Cost RT: MSE-hypothesized model = 1.03 (1.06), 95% CI [0.59 1.98], median MSE of permuted models = 1.01 (1.02), p = 0.65).

We also tested for a dissociation in our models, which included internetwork connectivity with the d-DMN and FPCN-A. In partial support of our third hypothesis we observed that the *resting-state* functional connectivity between FPCN-A, CON, DAN, and dorsal-DMN did not significantly predict median response time, but did predict mean accuracy of switch-costs better than a null distribution (Switch-Cost RT: MSE-hypothesized model = 0.91 (0.83), 95% CI [0.66 2.25], median MSE of permuted models = 1.09 (1.20), p = 0.03, Switch-Cost Accuracy: MSE-hypothesized model = 0.87 (0.76), 95% CI [0.64 1.78], median MSE of permuted models = 1.05 (1.10), p = 0.005). Similarly, we found that resting-state connectivity between FPCN-A, CON, DAN, and dorsal-DMN significantly predicted median response time but did not significantly predict median accuracy of inhibition-costs better than a null distribution (Inhibition-Cost RT: MSE-hypothesized model = 0.83 (0.67), 95% CI [0.53 2.22], median MSE of permuted models = 1.11 (1.23), p = 0.005, (Inhibition-Cost Accuracy: MSE-hypothesized model = 1.05 (1.10), 95% CI [0.58 1.83], median MSE of permuted models = 1.00 (1.01), p = 0.93).

We also observed that *task-based* connectivity between FPCN-A, CON, DAN and dorsal-DMN did significantly predict median response time, but not mean accuracy or median response time of switch costs and inhibition costs better than null distributions (Switch-Cost RT: MSE-hypothesized model = 0.87 (0.76), 95% CI [0.64 2.12], median MSE of permuted models = 1.15 (1.32), p = 0.005; Switch-Cost Accuracy: MSE-hypothesized model = 0.95 (0.90), 95% CI [0.69 1.91], median MSE of permuted models = 1.18 (1.39), p = 0.10; Inhibition-Cost Accuracy: MSE-hypothesized model = 0.69 (0.48), 95% CI [0.48 4.52], median MSE of permuted models = 1.43 (2.04), p = 0.03; Inhibition-Cost RT: (MSE-hypothesized model = 1.03 (1.06), 95% CI [0.57 2.00], median MSE of permuted models = 1.01 (1.02), p = 0.68).

### 3.4 Hypothesis 4: Correlations between cognitive networks during rest and task performance

Finally, we examined the correlation structure of CC networks at *rest* and during the Navon switching and Stroop inhibition *tasks* as our fourth hypothesis. We observed partial support for hypothesis 4a that connectivity between FPCN-B and dorsal-DMN at rest were negatively correlated (t (41) = -5.85, p *<*0.0001) and were positively correlated during the Stroop inhibition *task* (t (20) = 3.01, p =0.007) but not during the Navon Switching *task* (t (28) = 2.50, p = 0.02). We also observed partial support for hypothesis 4b. At *rest*, connectivity between FPCN-B and CON were not correlated (t (41) = 0.50, p = 0.62), but FPCN-B and DAN were positively correlated (t (41) = 9.23, p *<*0.0001). Further support for hypothesis 4b was observed when subjects performed the Navon switching *task*. We observed positive correlations in connectivity between FPCN-B, CON (t (28) = 8.24, p *<*0.0001), DAN (t (28) = 19.82, p *<*0.0001), and during the Stroop inhibition *task* when we observed positive correlations in connectivity between FPCN-B and CON (t (20) = 8.52, p *<*0.0001), DAN (t (20) = 18.68, p *<*0.0001). In partial support of our final sub hypothesis 4c, we observed that functional connectivity measured at *rest* was negatively correlated between dorsal-DMN and CON (t (20) = -9.07, p *<*0.0001), and DAN (t (20) = -10.52, p *<*0.0001). Interestingly, we saw that connectivity between dorsal-DMN and DAN remained negatively correlated during the Navon switching *task* (t (20) = -4.99, p *<*0.0001) and was not significant during the Stroop inhibition *task* (t (20) = -1.66, p = 0.11). In contrast, the inter-network connectivity of the dorsal-DAN and CON was positively correlated during the Navon switching *task* (t (20) = 4.01, p = 0.0004) and Stroop inhibition *task* (t (20) = 3.50, p *<*0.002) (see Figure 4).

**Fig. 4.**
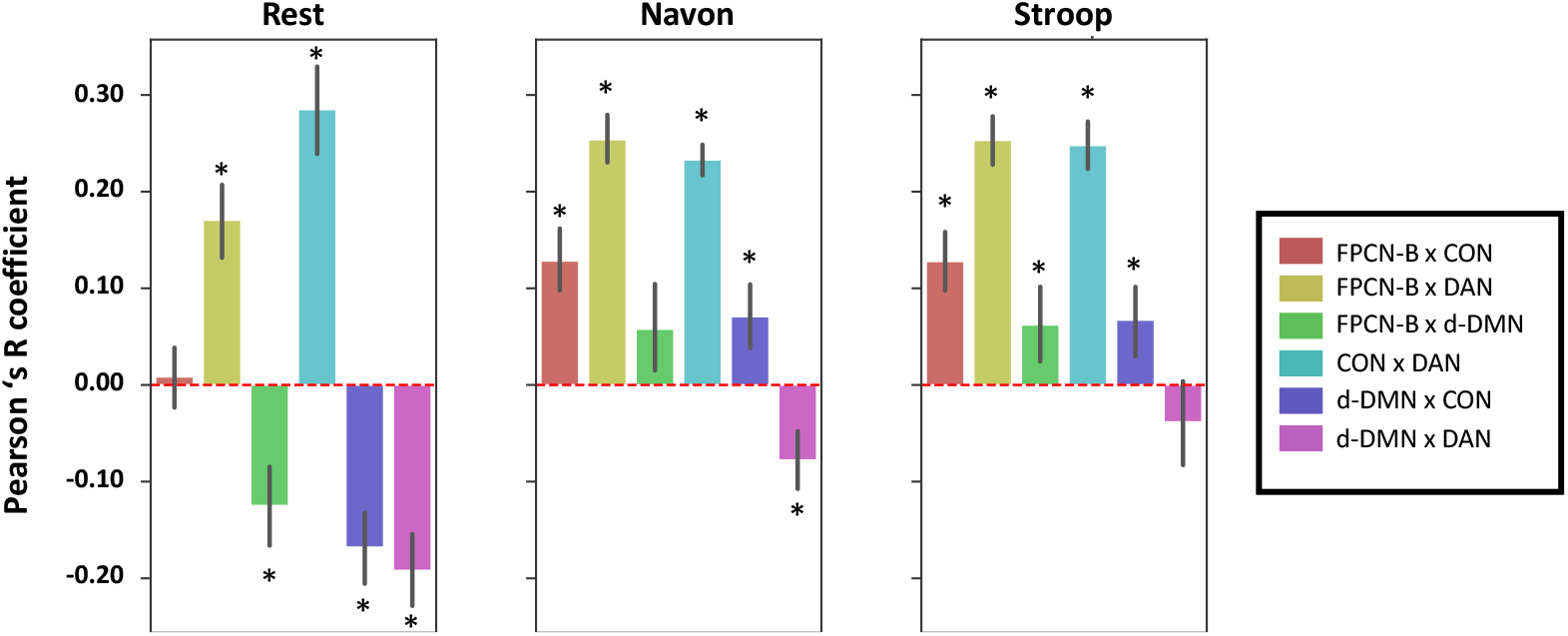
Bar chart displaying Pearson’s R correlations displaying pairwise inter-network connectivity from individuals. Error bars are estimated using 95% confidence intervals from the 10k OOB bootstrapping process. Depiction of Pearson’s R correlation for resting-state, Navon switching, and Stroop inhibition tasks.

### 3.5 Exploratory analysis of FPCN-A and CC correlation structure

To further understand the relationship between CC functional networks, we examined connectivity between FPCN-A, DAN, CON, and dorsal-DMN at *rest* and during *tasks*. During *rest*, we observed that connectivity between FPCN-A and CON (t (41) = -9.32, p *<*0.001) and DAN (t (41) = -8.35, p *<*0.001) were significantly negatively correlated and that connectivity between FPCN-A and the dorsal DMN were significantly positively correlated (t (41) = 13.96, p *<*0.001). During the Navon switching *task*, average connectivity between high and low conditions and between FPCN-A and CON were significantly positively correlated (t (28) = 2.86, p = 0.008) but not significantly correlated with DAN (t (28) = -1.25, p = 0.22), and was positively correlated with dorsal-DMN (t (28) = 12.59, p *<*0.001). Additionally, during the Stroop inhibition *task*, we observed a significant positive correlation of connectivity between FPCN-A and CON (t (20) =2.29, p = 0.03) and between FPCN-A and dorsal-DMN (t (20) = 18.85, p *<*0.001) but not DAN (t (20) = 0.72, p = 0.48).

## 4 Discussion

The current study investigated the relationships between the individually estimated FPCN, DMN, CON, and DAN and how these relationships predicted CC (i.e., switching and inhibiting) behaviors. We first tested whether restingstate and task-based functional connectivity between the FPCN-B, DAN, and CON significantly predicted switch- and inhibition-costs. Consistent with our first hypothesis, we observed that inter-network connectivity measured at rest and during task performance predicted response time and accuracy on a switching task and response time on a inhibition task. Supporting our second hypothesis, we observed that resting-state connectivity between the FPCN-B, CON, DAN, and dorsal-DMN predicted switching but not inhibition in individuals, while task-based connectivity predicting accurate switching and inhibition, but not response time. Partially consistent with our third hypothesis that FPCN-A and FPCN-B were responsible for dissociable behaviors, we observed at rest that connectivity between FPCN-A, DAN, and CON predicted switching accuracy and inhibiting response time. In contrast, this connectivity during tasks did not predict inhibiting or switching performance. Further, in support of our third hypothesis, we observed that during rest, connectivity between FPCN-A, DAN, CON, and dorsal-DMN predicted accurate switching and inhibiting accuracy, while during task performance, this connectivity predicted only switching response time, likely due to the inclusion of the dorsal-DMN. Finally, we observed support for our fourth and final hypothesis that FPCN-B and dorsal-DMN were negatively coupled at rest and positively coupled during the Stroop inhibition task. Further, in partial support of our fourth hypothesis, we observed that connectivity between FPCN-B and the CON was positively correlated during the Navon Switching and Stroop inhibition task. In contrast, connectivity between the FPCN-B and DAN was positively correlated at rest and during the Navon switching and Stroop inhibition tasks. We observed partial support for our final hypothesis, which indicated that the dorsal-DMN, CON, and DAN were negatively coupled at rest but were still negatively coupled with the DAN during the Navon switching task and positively coupled with the CON during the Navon switching and Stroop inhibition task.

We observed that inter-network connectivity between the FPCN-B, CON, and DAN was important in predicting switch-cost response time and inhibition-cost response time and accuracy. This result suggests that the relationships between these networks these networks may be interacting with one another to produce timely and accurate cognitive switching and inhibiting. Specifically, that the interactions between these networks are associated with effort required to switch and inhibit cognitive mechanisms. However, we did not observe support for these networks in predicting timely switching in individuals across task and rest data, indicating differences in functional dynamics between task and rest data [12]. While contrary to our hypothesis, our findings may indicate that the extra time taken to switch task sets may not meaningfully rely upon the FPCN-B, CON, and DAN. Instead, it could be that switching relies upon a different combination of these networks and others, such as the inclusion of the dorsal-DMN which may aide in updating information from previous trials [106, 107]. Interestingly, when we examined the relationship between our omnibus model, including the inter-network connectivity of FPCN-B, DAN, CON, and dorsal-DMN, we observed that the task-based inter-network connectivity of dorsal-DMN and DAN was predictive of accurate and timely switching and inhibiting behaviors in individuals. This result may suggest that the anti-correlation and non-correlated relationships seen during cognitive switching and inhibiting (See Figure4) tasks play distinct and dynamic roles [107]. Although previously detected within groups of healthy subjects [107], our results suggest that the relationships between these networks are dependent on the task at hand for individuals.

Findings from testing our second hypothesis suggest that, at rest, interactions among FPCN-B, CON, DAN, and dorsal-DMN provide baseline neural activity that is crucial for accurate and timely task switching. This result suggests that the dorsal-DMN’s role in aiding CC networks to update working memory based on contextual information may be related to CC networks at rest [107]. Additionally, the finding that connectivity between FPCN-B, DAN, CON, and dorsal-DMN during task performance predicts accurate switching and timely inhibiting suggests that the dynamic activity between these networks is vital for a subject’s ability to switch task set rules during task performance accurately but is also important for updating performance (possibly increasing response hesitancy) due to previous trial performance or task completion heuristics. However, future studies would need to directly test this hypothesis to confirm these ideas.

The findings from our first and second hypotheses suggest that the relationship between FPCN-B, dorsal-DMN, CON, and DAN networks are vital for inhibiting prepotent stimuli and switching/ working memory task-set updating. Individual connectivity derived from tasks and during rest shows that the anti-correlated dorsal-DMN and DAN were important factors for that predicted accurate and timely inhibiting and set switching. Indeed, at the group level, these relationships associated with working memory performance have been previously established [108, 109]. However, to our knowledge, we are the first to observe these relationships within individuals, and that task and rest functional connectivity can predict inhibiting and set-shifting. These observations provide further evidence that these networks likely work together to promote accurate and timely inhibition of prepotent responses and set switching and updating. That we observed this relationship within individuals confirms previous work by Dixon and colleagues (2017) who showed that the functional relationship between the FPCN, dorsal-DMN and DAN were dependent on the task at hand [107]. However, this study did not seek to establish a relationship between inter-network connectivity and behaviors. Furthermore, it places the relationship between the dorsal-DMN and the DAN as important for switching and inhibiting even in a model with other canonically task positive relationships. This suggests that the segregation between the dorsal-DMN during task switching task performance is associated with individual switching performance. Further, the segregation between these two networks at rest is also important for accurately inhibiting automated responses.

Our findings also support our third hypothesis, that connectivity between FPCN-A and other CC networks is less critical for accurate and timely switching and inhibiting. Indeed, previous studies have suggested that individuals performing a task need to predict future events and that cognitive branching may require more anterior nodes of the DLPFC, similar to those found in FPCN-A [36, 39]. Interestingly, we found that FPCN-A is important for switching and inhibiting, but only during rest.

Our findings from our second and third hypotheses suggest a possible rest/task division between FPCN-A and B. Specifically, we show that during rest, FPCN-A and B interact with other CC networks and that these interactions are important for timely and accurate switching and inhibition. These results and our exploratory analysis suggest that it may be necessary for FPCN-A to disengage with other CC networks while performing a task. On the face of it, these results seem to contradict previous studies which compare task and rest functional connectivity and find that they are broadly similar [110]. However, previous research (including with individually estimated connectivity) also suggests that these subtle differences between task and rest connectivity are important for task execution [28, 29, 110–112].Specifically, one previous study showed that within and between network differences in functional connectivity were present in task-based compared to resting-state data and that inter-network connectivity increased during task performance compared to rest, regardless of the task, and primarily between different control regions such as the FPCN, CON, and DAN, and DMN. Furthermore, they also observed that during task performance, regions such as the FPCN displayed increased between network connectivity and decreased within network connectivity compared to resting state functional connectivity [28]. The current study shows that individually-estimated functional networks provide important distinctions between rest and task performance for switching and inhibiting behaviors. Again, these results confirm previous studies showing that task-rule switching and inhibiting paradigms recruit differential nodes of the FPCN, most notably along the rostral-caudal axis of the DLFPC [36, 37, 39]. These results suggest that CC in individuals likely follows similar rostral-caudal axis information processing. Our results are also consistent with a previous study examining response inhibition in healthy adults after deploying a factorial design of another type of task measures inhibition, the go/no-go task, which revealed stronger negative functional connectivity between the inferior frontal gyrus (a node in the CON), the precuneus, and posterior cingulate gyrus (both nodes of the DMN) during response inhibiting trials [113], interestingly, these are both nodes within the canonical DMN. The current study shows that connectivity between the CON and dorsal-DMN, FPCN-A and B, and the DAN is critical in determining accurate and timely responses to tasks requiring inhibiting automated responses and switching task-rule sets.

The results of previous and the current study suggest that for tasks requiring CC, successful inhibiting and task set switching require both the FPCN-A and FPCN-B, albeit differently during rest and when performing a task. Specifically, FPCN-A may be involved in pre-trial planning (i.e., cognitive branching and creation of sub-goals) and may need to be disengaged during switching and inhibiting execution. In contrast, FPCN-B may be engaged in momentto-moment trial execution and post-trial re-calibration within individuals [28]. Supporting these proposed functions are studies using functional activity maps between subjects and observed differences in activation between regions of the lateral prefrontal cortex (LPFC)[23, 32, 36, 37, 40], while caudal regions were associated with the integration of sensory information from areas such as the frontal eye fields (part of DAN) and motor processing regions [32]. Importantly, this functional dissociation in activity patterns and behavior also maps to the functional dissociation of FPCN-A and FPCN-B. Furthermore, Nee and D’Esposito (2016, 2017) showed that rostral regions of the bilateral LFPC were also associated with predicting future process demands and the timing of a response, suggesting a more top-down role for CC facilitated by regions like those encompassing FPCN-A [37, 39]. Additionally, caudal areas of the bilateral LPFC were related to current processing demands or moment-to-moment processing required for a given task trial and are related to bottom-up CC and map onto FPCN-B. Research also suggests that an anterior-posterior divide between the LPFC exists such that the anterior portion of the LPFC is associated with forming sub-goals for a given task. In contrast, the posterior portion of the LPFC is necessary for sensory processing integration and control; both aspects of control would seem necessary for successful and timely performance of the Navon switching and Stroop inhibition [26, 32]. These studies suggest complementary roles for the FPCN depending on topographical location [36, 37, 39]. Our results indicate that the relationships between individually-estimated functional networks at rest and during CC tasks are essential in different ways for individual task performance.

The results addressing our final hypothesis concerning the dynamics between networks associated with CC are consistent with previous findings. Our findings show that individually measured connectivity between FPCNB and dorsal-DMN display tight coupling when performing tasks and are anti-correlated at rest. Here we show that this relationship exists for tasks associated with CC (task-set switching and inhibiting automated responses in individualized networks) [10, 35]. The relationship between FPCN-B and dorsal-DMN suggests a positive interaction between the two networks where dorsal-DMN may support CC. Specifically, it may be that dorsal-DMN aides in switching and inhibiting cognition by retrieving and binding contextual details to simulate future events and thus correcting trial-level behaviors [63, 107**?**]. To our knowledge, this is the first study showing this relationship within individuals and provides further evidence of the importance of this network to task-set switching and inhibiting automated responses.

The functional networks investigated in this study and our ability to predict switching and inhibiting behaviors using the relationships between these CC networks validate the importance of these networks to basic CC behaviors. Importantly, we also observed that individual differences in task performance measuring switching and inhibiting can be predicted by individually estimated functional relationships between functional networks in the human brain.

## 5 Limitations and Future Directions

We note some limitations to our study. First, studies suggest the most reliable estimates of intrinsic connectivity are obtained from approximately 30 minutes of resting-state data [114]. However, our connectivity matrices were derived from 8 to 16 minutes per subject. A second limitation is that our task design consisted of equal blocks to maximize the ability to analyze connectivity, making it challenging to extract trial-level data. Future studies should also use trial-level regression to examine how these networks contribute to trial-level temporal dynamics within individuals. Another limitation to our study includes using complex nonlinear kernels for the SVR model. While nonlinear kernels increase the power of SVR models to detect potentially complex associations with outcomes, using these kernels diminishes our ability to interpret the individual features. Specifically, because of our model optimizer selection, there are still unanswered questions about the unique and shared variance in predicting inhibiting and switching behaviors as the nonlinear nature of the optimizers do not provide interpretable weights associated with predicting switching and inhibiting. Studies in the future will need to adopt more sophisticated predictive modeling schemes to answer these questions directly, preferably those with interpretable weights from the modeling fit. Finally, throughout this article we refer to the terms co-vary and between network connectivity. We do not assume that these networks are directly interacting with one another but that the statistical relationship between these networks may be a meaningful measure of individual variance related to behavior. As all functional connectivity studies are correlative by nature, the field could benefit from considering structural constraints and mediators between regions.

## 6 Conclusions

In this study we provide evidence that individualized inter-network connectivity can predict CC behaviors. This study provides important groundwork for understanding the relationship between functional networks within individuals and their relationship to individual CC abilities. We show that individual differences between functional networks are predictive of an individual’s ability to perform two of three commonly associated CC behaviors (switching and inhibition) [6] depending on whether an individual is at rest of engaged in a cognitive task.We also provide evidence that CC networks likely work in concert with one another within individuals to bring about accurate and timely switching and inhibiting behaviors. In sum, these findings suggest that individual differences in between-network functional connectivity amongst the FPCN and other CC-associated networks can predict an individual’s ability to accurately and timely inhibit automated responses and switch task rules. Studies such as ours will be crucial to understanding individuals with deficits related to CC and may begin to shed light on individual differences in the dynamics between functional networks and how those dynamics facilitate performance on cognitive tasks. However, it is important to acknowledge that the temporal aspect of how these networks emerge within individuals is likely key to fully understanding the function of CC networks, thereby requiring more sophisticated temporal assays of CC networks in the human brain combined with the resolution advantages of fMRI. Although, the approach taken in this study using individually estimated inter-network functional connectivity to predict individual CC abilities should still promote new studies in clinical populations associated with executive control deficits such as Attention-Deficit Hyperactivity Disorder (ADHD) and other psychiatric and neurological conditions.

## Acknowledgments

We would like to thank all participants in this study for their dedication to furthering our knowledge of basic cognitive functions.

