## Supplemental Tables and Figures for "Individual-level Functional Connectivity Predicts Cognitive Control Efficiency"

### Individual-level Functional Connectivity Predicts Cognitive Control Efficiency: Supplementary Information and Tables

#### 1 Supplement

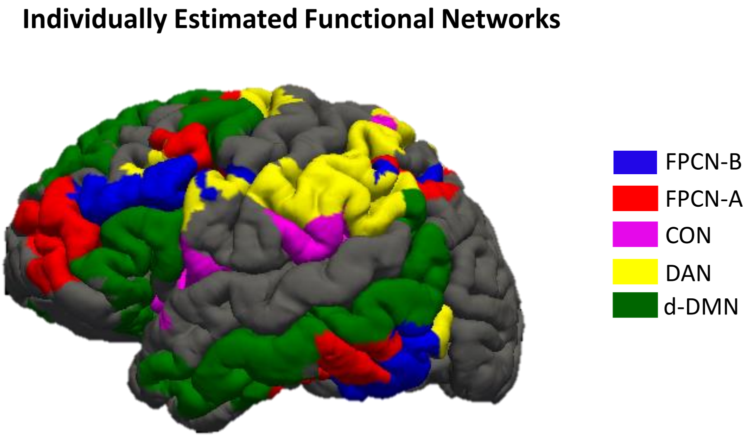


**Fig. 1** Showing all individually estimated functional networks overlaid onto an individual's surface reconstruction.

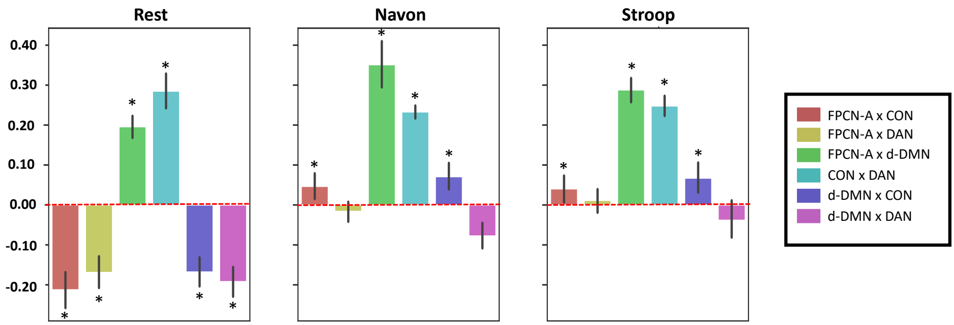


**Fig. 2** Pearson's R correlations displaying pairwise inter-network connectivity from individuals. Error bars are estimated using 95% confidence intervals from 10k OOB bootstrapping process. Depiction of Pearson's R correlation for resting-state, Navon switching, and Stroop inhibition tasks.

| Parameter | Value(s) |
|---|---|
| Kernel | Radial-Basis Function, Sigmoid, Linear |
| Epsilon (if RBF Kernel) | 0.00001, 0.0001, 0.001, 0.01, 0.1, 1, 10, 100, 1000 |
| Cost | .001,.01,.1,1,10,100,1000,1000 |
| Gamma | .000001,.00001,.0001,.001,.01,.1,1,10 |

**Table 1** List of Possible SVR hyperparamters used in Grid-Search CV

| Parameter | Values | Even splits |
|---|---|---|
| Number of Trees | 100 - 1000 | 10 |
| Number of features at split | auto, square-root | |
| Maximum number of levels in tree | 10-110 | 11 |
| Minimum number of samples required to split node | 2,5,10 | |
| Minimum number of samples required at each leaf | 1,2,4 | |

**Table 2** List of possible Random Forest Regression parameters
